# Literature-based predictions of treatments for genetic disease pathology

**DOI:** 10.1101/2022.09.08.506253

**Authors:** Cole A. Deisseroth, Won-Seok Lee, Ji-Yoen Kim, Hyun-Hwan Jeong, Julia Wang, Huda Y. Zoghbi, Zhandong Liu

## Abstract

Identifying genetic modifiers of disease-causing genes can guide drug discovery for treatment of genetic disorders, but selecting promising drugs to test requires extensive and up-to-date knowledge of drug-gene and gene-gene relationships. To address this challenge, we present PARsing ModifiErS via Abstract aNnotations (PARMESAN), a computational tool that searches PubMed for information on these relationships, and assembles them into one knowledgebase. PARMESAN then hypothesizes on undiscovered drug-gene relationships, assigning an evidence-based score to each hypothesis. We compare PARMESAN’s drug-gene hypotheses to all of the drug-gene relationships displayed by DrugBank, and see a strong correlation between the prediction score and the predictive accuracy—such that predictions scoring above 10 are 11 times more likely to be correct than incorrect. This publicly available tool provides an automated way to prioritize drug screens to target the most-promising drugs to test, thereby saving time and resources in the development of therapeutics for genetic disorders.

## Main text

The Undiagnosed Diseases Network has aided in the discovery of many Mendelian disorders, but each one requires its own treatment. Due to the large number of disorders and the small number of individuals affected by each one, searching for treatments is impractical without proper guidance. Identifying therapeutics for a genetic disorder quickly and systematically requires a means of deciding which therapeutics to test first. Making this decision requires an understanding of the molecular pathway and the roles of possible therapeutics in that pathway. Existing databases of gene-gene and drug-gene relationships can aid in the exploration of these molecular pathways, but may not be enough to inform hypotheses on therapeutic effects.

Protein-protein interaction databases, such as BioGRID^1^, STRING^2^, BioPlex^3^, and Reactome^4^, are large enough to enable network analyses, but they do not provide directionality—whether one protein increases or decreases the activity of another by interacting with it.

There exist substantial gene-gene relationship databases such as the Kyoto Encyclopedia of Genes and Genomes (KEGG)^5^, which provides more than 50,000 of these relationships, across more than 5,000 genes, but that means that there is less than a 25% chance that this database will have such information for a given gene. Other genetic modifier databases, such as NeuroGeM^6^ and PhenoModifier^7^, and drug databases such as DrugBank^8^, are manually curated, and there is no sustainable way to update them with the rapidly growing body of scientific literature.

There currently exist tools that show promise in prioritizing treatments for rare disorders, such as mediKanren^9^, which aggregates publicly available knowledgebases that link biomedical concepts, and broadly hypothesizes on links between biological entities, including possible disease therapeutics. To our knowledge, however, there are no publicly available tools aimed at deriving drug and genetic modifier predictions directly from the primary literature, which is the first place that newly discovered relationships will be displayed—and for many discoveries, possibly the only place.

To aid in the investigation of therapeutics for rare disorders, we present PARMESAN (PARsing ModifiErS via Abstract aNnotations), an automated literature search tool that scans through every abstract in PubMed for descriptions of gene-gene and drug-gene relationships, and encodes these relationships into one database. PARMESAN then uses this database to hypothesize on the effect a given drug or protein has on a given disease gene. We test the validity of PARMESAN’s drug hypotheses by comparing them to all of the drug-gene relationships presented in DrugBank.

While it aids directly in the search for disease therapeutics, PARMESAN can also facilitate the discovery of gene-gene relationships, which improves our understanding of molecular pathways, and further boosts the identification of promising therapeutics. We compare PARMESAN’s gene-gene relationship hypotheses to the gene-gene relationships available in KEGG, and to genetic modifier screens of two disease genes encoding proteins that play important roles in neurodegeneration or neurodevelopment: *Ataxin 1* (*ATXN1*), a protein whose unstable CAG trinucleotide repeat expansion causes neurodegeneration in Spinocerebellar Ataxia Type 1^10,11^; and *Microtubule-Associated Protein Tau* (*TAU*), a protein that has roles in Alzheimer’s Disease, frontotemporal dementia, and Parkinson’s Disease^12–14^.

By comparing PARMESAN’s predictions to known gene-gene and drug-gene relationships, we demonstrate its ability to generate valid hypotheses from the primary literature, and its potential in prioritizing the best hypotheses to investigate.

## Results

### PARMESAN builds a substantial knowledgebase of drug-gene and gene-gene relationships

PARMESAN processes abstracts and summarizes the modifier relationships that they describe in the following format: “Action Modifier Predicate Target”. The action is the way the modifier gene is being manipulated, and the predicate is the effect that manipulating the modifier has on the target gene. As an example, Monteiro et al published an article titled, “Pharmacological disruption of the MID1/_α_4 interaction reduces mutant Huntingtin levels in primary neuronal cultures”^15^. The modifier and target are MID1 and Huntingtin, respectively. The predicate—what MID1 is doing to Huntingtin—is “reduces”. And the action—what is being done to MID1 to achieve this effect on Huntingtin—is “disruption”. The resulting sentence is therefore “disruption MID1 reduces Huntingtin” (Figure 1).

**Figure 1:**
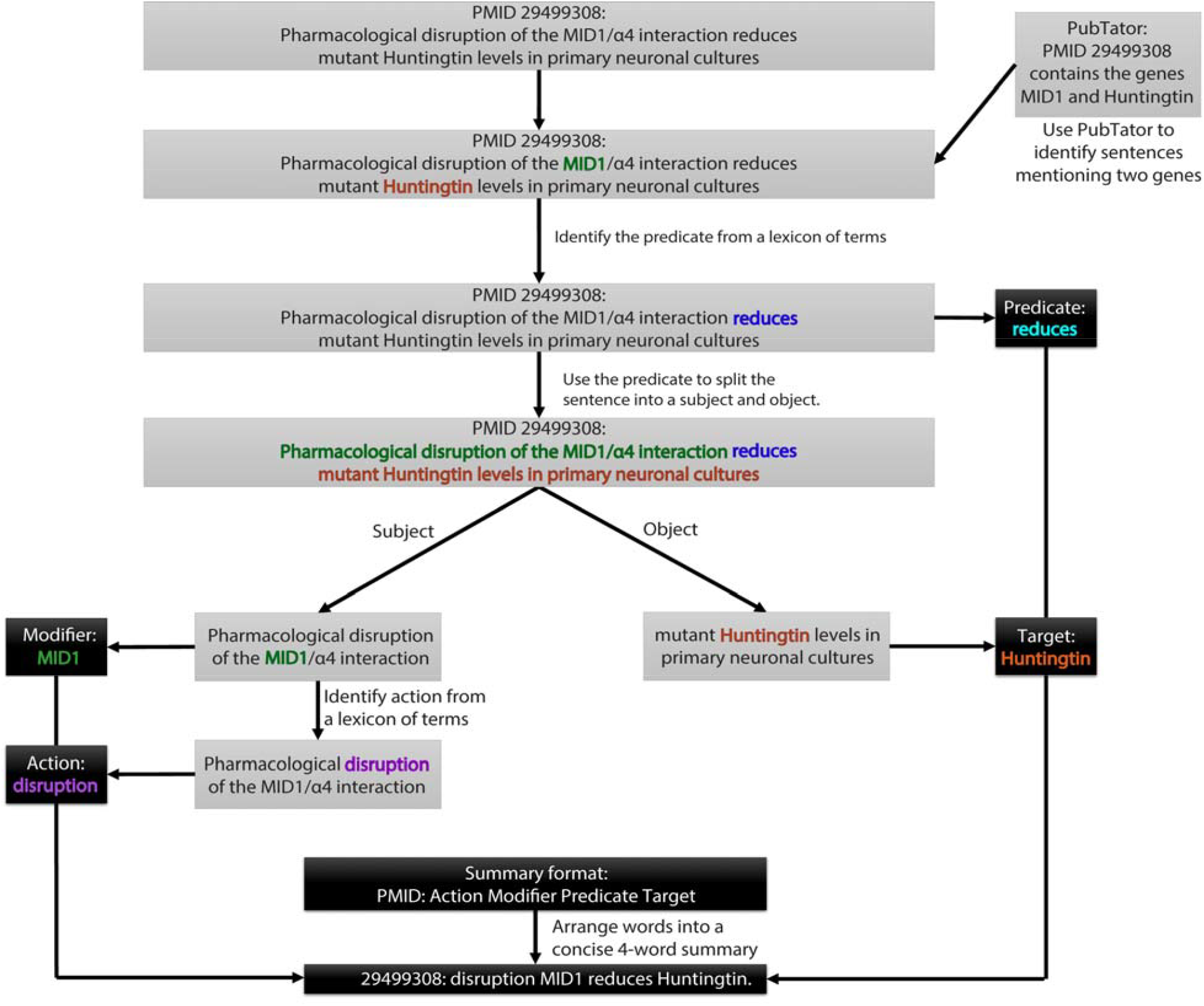
Summary sentence construction. PARMESAN constructs summary sentences from abstract sentences in the following format: Action Modifier Predicate Target, where the Predicate is the effect the modifier has on the target gene (such as increase or decrease), and the action is what is being done to the modifier to achieve this effect (whether the modifier should be increased or decreased to have the mentioned effect on the target). The Action is the only optional part of the summary sentence.

We gave PARMESAN a vocabulary of 70 positive predicates, 142 negative predicates, and 63 negative actions (Table 1). PARMESAN constructed a knowledgebase of known gene-gene relationships, which were positively (the modifier, which we will call a “positive regulator”, increases the level or activity of the target gene’s protein) or negatively directed (the modifier, a “negative regulator”, decreases the level or activity of the target gene’s protein). PARMESAN automatically identified hundreds of thousands of relationships among tens of thousands of genes (indexed by Entrez gene ID, Table 2).

**Table 1:** Vocabulary used by PARMESAN. A positive predicate is a word that suggests an increase in the expression or activity of the target gene. A negative predicate is a word that suggests a decrease in the expression or activity of the target gene. A negative action is a word that suggests decreased expression or activity of the modifier. Positive action words are not considered, because they do not add any information in the absence of a negative action. For example, “Gene A suppresses Gene B” has the same meaning as “Increasing Gene A suppresses Gene B”, as both indicate Gene A to be a negative regulator of Gene B.

|  | **After** |
| --- | --- |
| **Positive predicate** | accelerate, accelerated, accelerates, activate, activated, activates, allowed, allows, augment, augmentation, augmented, augments, coactivator, costimulate, elevate, elevated, elevates, elicit, elicited, elicits, enable, enables, enhance, enhanced, enhancement, enhancer, enhances, enhancing, exacerbate, exacerbated, exacerbates, expedited, facilitate, facilitated, facilitates, improve, improved, improvement, improves, induce, induced, -induced, induces, potentiate, potentiated, potentiates, promote, promoted, promoter, promotes, reactivate, reactivated, reactivation, restore, restored, restores, stabilize, stabilized, stimulate, stimulated, stimulates, strengthened, strengthening, strengthens, transactivated, transactivating, transactivation, upregulate, upregulated, upregulates |
| **Negative predicate** | ablated, abolish, abolished, abolishes, abolishing, abrogate, abrogated, abrogates, antagonised, antagonises, antagonize, antagonized, antagonizes, coactivated, coinhibition, compete, competes, counteract, counteracted, counteracting, counteracts, deactivated, deactivates, degrade, degraded, degrader, depressed, destabilize, destabilized, destroyed, destroying, diminish, diminished, diminishes, diminishing, disassembled, disassembles, displace, displaced, displacement, displaces, disrupt, disrupting, disruption, dissociate, dissociated, disturb, disturbances, disturbed, disturbing, disturbs, divert, downregulate, down-regulate, eliminate, eliminated, eliminates, hinder, hindered, hindering, hinders, impair, impaired, impairing, impairment, impairs, impede, impeded, impedes, inactivate, inactivated, inhibit, inhibited, inhibiting, inhibition, inhibitions, inhibitive, inhibitory, inhibits, interceptor, interfere, interfered, interference, interferes, interfering, interrupt, interrupted, interrupting, interruption, interrupts, negated, negates, neutralize, neutralized, neutralizes, nullify, obliterated, obstructive, oppose, opposed, opposes, outcompeted, perturbation, perturbed, prevent, prevented, preventing, prevention, prevents, prohibited, prohibiting, reduce, reduced, reduces, reducing, repress, repressed, represses, repressing, repressive, restrict, restricting, restricts, revert, reverted, reverting, suppress, suppressed, suppresses, suppressing, suppression, suppressive, terminated, terminates, transrepresses, transrepression, undermined, unstimulated, weaken, weakened, weakening, weakens |
| **Negative action** | abolished, abrogated, absence, antibody, blockage, blocked, blocking, decrease, decreased, decreasing, deficiency, deficient, deleted, deleting, deletion, depleted, depleting, depletion, diminished, disruption, downregulate, down-regulate, downregulated, down-regulated, downregulates, downregulating, down-regulating, downregulation, down-regulation, eliminated, impaired, inhibit, inhibited, inhibiting, inhibition, inhibitor, inhibitors, inhibitory, inhibits, interfere, interference, interfering, knock, knockdown, knock-down, knocked, knocking, knockout, knock-out, loss, neutralizing, prevent, prevented, reduce, reduced, reduces, reducing, rnai, sirna, suppressed, suppresses, suppressing, suppression |

**Table 2:** Statistics on the initial knowledgebase constructed. PARMESAN automatically found hundreds of thousands of modifier relationships in the medical literature.

|  | Articles | Modifiers | Targets | Relationships |
| --- | --- | --- | --- | --- |
| Positive drugs | 118,922 | 9,045 | 21,580 | 136,055 |
| Positive drug predictions | 425,881 | 13,374 | 37,575 | 13,472,902 |
| Negative drugs | 99,694 | 10,747 | 18,140 | 131,517 |
| Negative drug predictions | 422,441 | 13,395 | 37,245 | 15,191,901 |
| Positive genetic modifiers | 260,077 | 39,794 | 34,935 | 487,274 |
| Positive genetic modifier predictions | 341,486 | 47,703 | 40,293 | 46,877,005 |
| Negative genetic modifiers | 170,467 | 35,545 | 29,997 | 335,692 |
| Negative genetic modifier predictions | 341,210 | 47,354 | 40,038 | 41,234,297 |

### PARMESAN’s extraction confidence scores associate strongly with the accuracy of extracted relationships

To test PARMESAN’s accuracy, we took a random sample of 25 extracted gene-gene relationships, and manually checked the articles supporting each relationship. There were three possible outcomes: 1. Correct—the above finding that “disruption MID1 reduces Huntingtin” was correctly extracted from the source article. 2. Misdirected—the finding that “p38 induces HDAC4” (Supplementary table 1) extracted from Zhou et al accurately reflected that p38 modifies the activity of HDAC4, but the article suggests that p38 actually induces the degradation of HDAC40, indicating a negative, rather than a positive effect^16^. 3. Incorrect—The modifier is not indicated to modify the activity of the target in the source article. As an example, the finding from Mukhopadhyay et al that “HeyL reduced TrkC” (Supplementary table 1) was not depicted in the source article— the target, TrkC, was being used to mark cells that were being reduced in number, but the protein itself was not indicated to be reduced in level, effect, or activity^17^.

Among the random gene-gene relationships, 60% were found to be correct (Figure 2). We performed the same test on a random sample of 25 drug-gene relationships extracted by PARMESAN, and 68% were correct.

**Figure 2:**
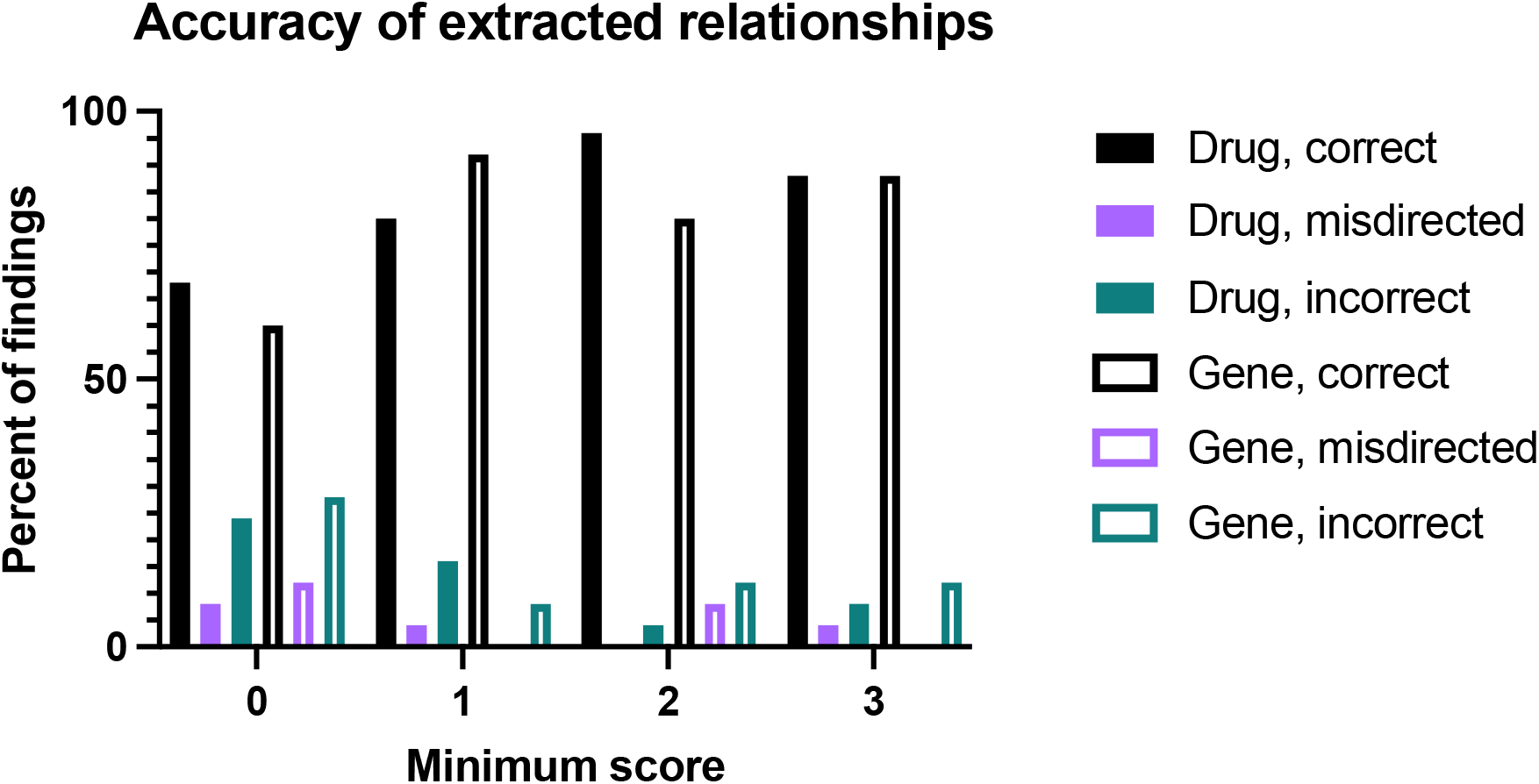
Accuracy of extracted modifier relationships: In eight different trials, we randomly selected 25 relationships extracted by PARMESAN, and checked the supporting articles to see whether at least one of them truly signified the relationship. The black columns represent the percent of relationships that were confirmed—with correct directionality—in at least one of the supporting articles—for example, PARMESAN states that Gene A negatively regulates Gene B, and a supporting article indicates that Gene A negatively regulates Gene B. The purple columns represent the number of relationships that were confirmed by at least one supporting article, but never with correct directionality—for example, PARMESAN states that Gene A negatively regulates Gene B, and the supporting articles indicate that Gene A positively regulates Gene B. The green columns represent the number of relationships that were not confirmed by any of the supporting articles. For each minimum score, we randomly select exclusively among the relationships that were given a confidence score whose absolute value was above that score. PARMESAN’s accuracy significantly improves when relationships are limited to those with a score above 3 (or below -3), suggesting that these scores are a promising measure of confidence in an automatically extracted relationship.

PARMESAN’s accuracy improved when we limited its findings to those with high scores. For n = 1, 2, and 3, we performed the same accuracy test on 25 random gene-gene and 25 random drug-gene relationships that were given a confidence score above n. At n = 3, the accuracy increased from 60% to 88% (p=0.004293) for gene-gene relationships and from 68% to 88% (p=0.0051458) for drug-gene relationships (Figure 2). This filter leads to a statistically significant improvement in accuracy for both gene-gene and drug-gene relationships.

### PARMESAN’s higher-scoring predictions are more likely to match extracted relationships

We ran PARMESAN’s gene-gene relationship predictor (Figure 3a) using only articles published before January 1, 2010. We then compared the predictions to relationships extracted from articles published before 2010 and before 2020 (Figure 3b, Supplementary table 13). We only used extractions with an extraction confidence score above 1.0. We consider a prediction “validated” if PARMESAN extracted the predicted relationship from the literature, with the same proposed directionality (both agree on whether Gene A increases or decreases the activity of Gene B). 0.2% of PARMESAN’s pre-2010 predictions were validated by 2020. However, requiring higher prediction scores increased the validation rate, and 68% of the predictions scoring above 20 were validated by 2020 (p value undetectably low).

**Figure 3:**
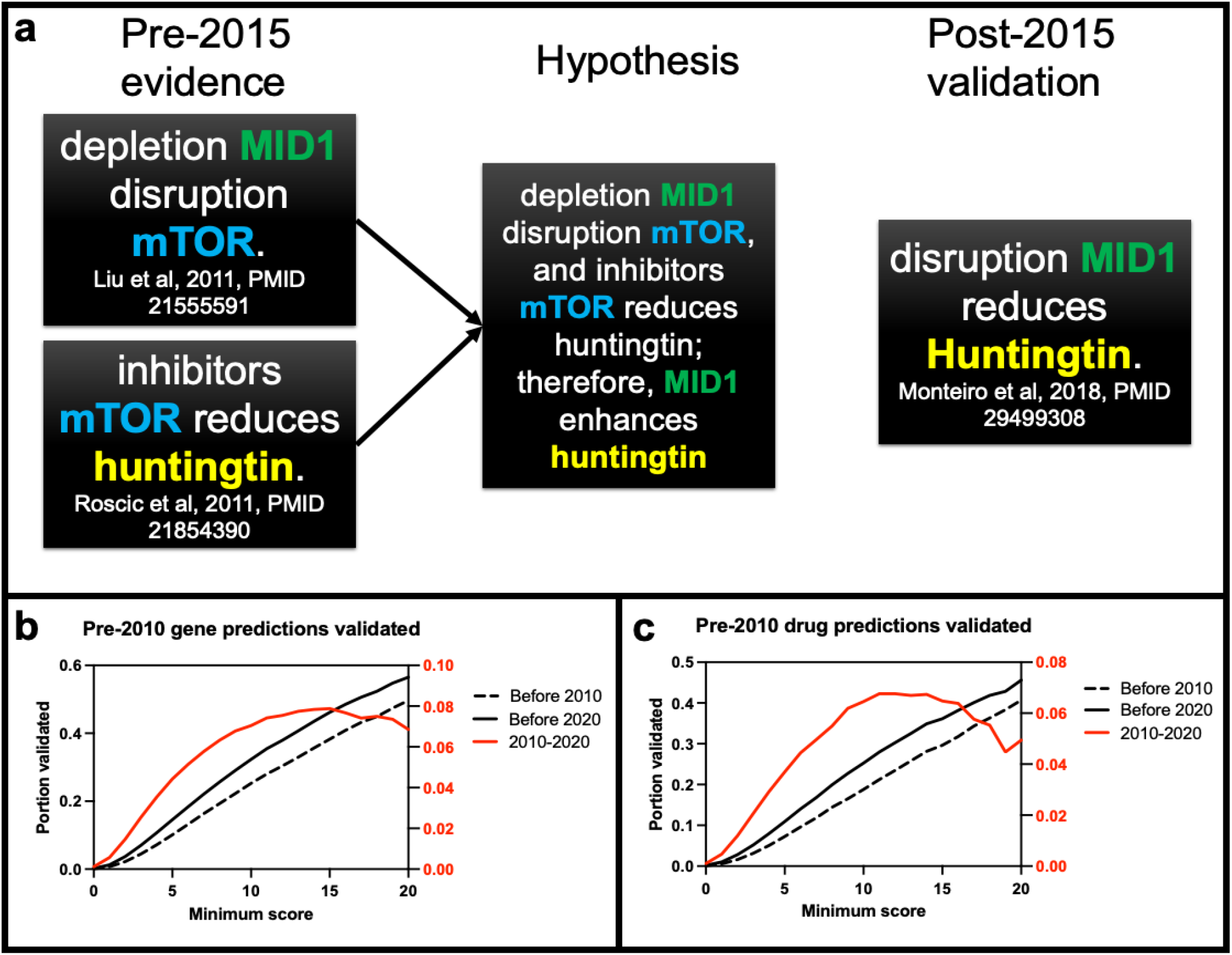
Validation of pre-2010 predictions over time. (**a**) We test PARMESAN’s long-term predictive capability with a time-capsule test, where PARMESAN makes predictions of modifiers using abstracts from before a given year, and we then compare those predictions to modifiers reported on after the given year. (**b**,**c**) We used PARMESAN to predict gene-gene (**b**) and drug-gene (**c**) relationships using only articles from before 2010, and show the fraction of those predictions that were consistent with relationships extracted before 2010 (black solid line) and before 2020 (black dashed line). The difference between these values (shown in red) represents the change in the fraction of predictions that were consistent with identified relationships over the decade after the predictions were made.

Unexpectedly, the rate of new validations (measured as the difference between the validation rates before 2010 and before 2020) peaked at 8% with a minimum score of 15, and fell to 7% with a minimum score of 20 (p < 0.005). This suggests that predictions with high confidence scores are more likely to be accurate, but less likely to yield a novel discovery.

We performed the same test with the pre-2010 drug-gene relationships (Figure 3c, Supplementary table 13), and saw a similar pattern: 0.2% of all predictions and 46% of predictions scoring above 20 were validated within the next 10 years (p value undetectably low), and the rate of novel validations peaked at 7% with a minimum score of 12 (p < 10^−5^).

### PARMESAN accurately predicts the effects of drugs on genes

We compare PARMESAN’s predicted drug-gene relationships to the drug-gene relationships described in DrugBank (Figure 4a, Supplementary table 14). We show that, when PARMESAN’s prediction confidence score is greater than 10, the predictions are 11 times more likely to be proven correct than incorrect. As modifiers are filtered based on their score, the correct predictions are significantly favored over the incorrect ones (p=0.0003).

**Figure 4:**
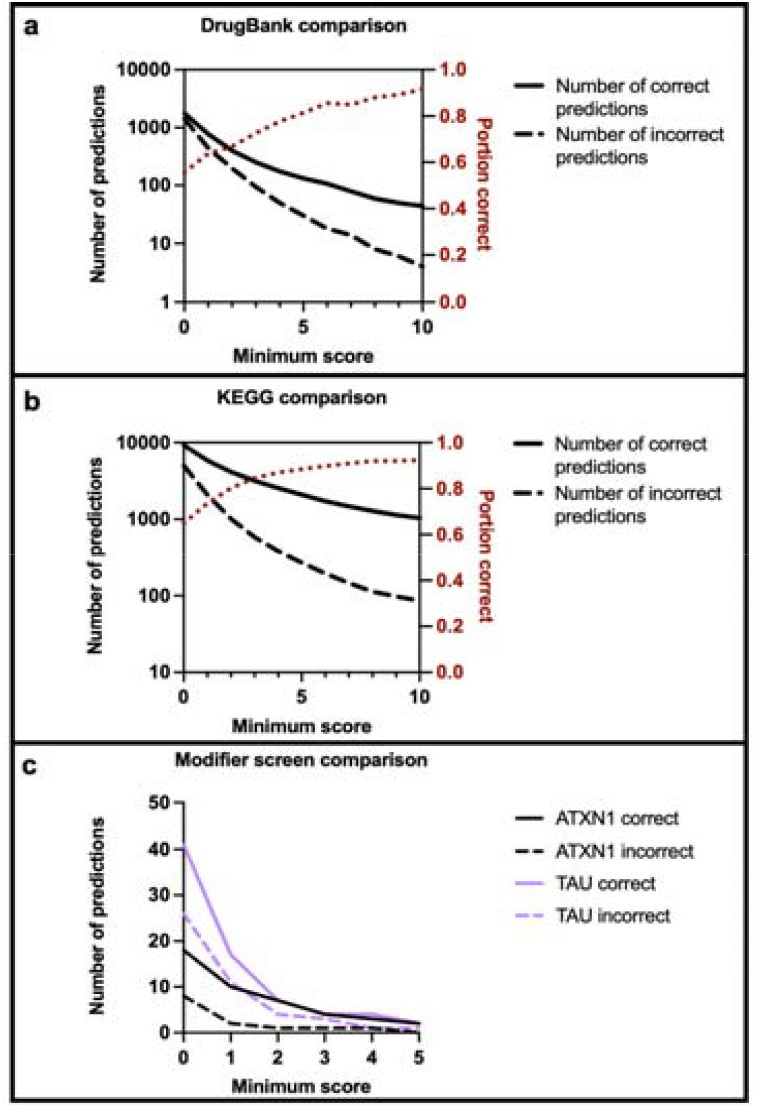
Comparison of PARMESAN’s gene-gene and drug-gene relationship predictions to experimentally validated relationships. We compared all of PARMESAN’s predictions to known drug-gene and gene-gene relationships. “Correct” means that the directionality predicted by PARMESAN (Whether Drug A positively or negatively regulates Gene B) matches that of the known relationship, and “Incorrect” means that PARMESAN predicted the opposite directionality (Drug A positively regulates Gene B, but PARMESAN predicted that Drug A negatively regulates Gene B). (**a**) The drug-gene relationship predictions were compared to the relationships presented in DrugBank, and predictions with scores at or above 10 are 11 times more likely to be proven right than wrong. (**b**) Gene-gene relationship predictions were compared to the gene-gene relationships presented in KEGG, and predictions with scores above 10 are 12 times more likely to be correct than incorrect. (**c**) We compared PARMESAN’s genetic modifier predictions for *ATXN1* and *TAU* to corresponding modifier screens, and the correct predictions outnumbered the incorrect ones.

### PARMESAN accurately predicts the effects of genes on other genes

We compare PARMESAN’s predicted gene-gene relationships to all of the gene-gene relationships described in KEGG^5^ (Figure 4b, Supplementary table 15). A minimum confidence score of 10 was associated with a correct:incorrect ratio of 12:1. Requiring a higher minimum score led to significant favoring of correct predictions (p<0.0001).

Lastly, we compare PARMESAN’s gene-gene relationship predictions to two *in vitro* disease gene modifier screens. One for genes whose knockdown decreases *ATXN1* levels^18^, and one for druggable genes whose knockdown increases or decreases endogenous and over-expressed *TAU* protein in a human cell line. PARMESAN’s correct predictions outnumber the incorrect ones, and filtering the predictions by score significantly favors the correct predictions in the *ATXN1* screen, but not the *TAU* screen (p=0.0003 and 0.5568, respectively).

## Discussion

Testing a drug for therapeutic effects in a given disorder is an expensive and time-consuming process, and it is essential to know which treatments are the most likely to work, especially for newly discovered genetic disorders for which no therapeutics have been developed. PARMESAN’s drug-predicting capability has the potential to save time and resources in the investigation of promising therapeutics for disorders caused by protein haploinsufficiency or toxic gain-of-function mutations.

The predictive power of this tool expands with time, as new publications emerge that report gene-gene and drug-gene relationships. In theory, the most effective way to sustainably collect information on gene-gene and drug-gene relationships would be to require researchers to upload new discoveries to a public database—just as sequencing and phenotypic data from studies can be mandatorily uploaded to the Database of Genotypes and Phenotypes (dbGaP). Until such a system exists for gene-gene and drug-gene relationships, automated literature curation may be the next best option.

PARMESAN can construct an entire knowledgebase automatically, accomplishing years of manual curation within hours, and can re-do the process every day to capture the most-recent publications. This knowledgebase contains hundreds of thousands of relationships, which are more than 50% accurate overall, and more than 80% accurate if they scored above 3.

The known relationships allowed PARMESAN to automatically predict unconfirmed ones, and the confidence scores are a strong indicator of predictive accuracy when compared to both extracted and database-derived relationships. Drug predictions with scores above 10 are 11 times more likely to be proven right than wrong. These drug predictions may therefore guide the development of therapeutics for diseases caused by heterozygous loss-of-function mutations (for which the goal would be to increase the activity of the healthy protein), or gain-of-function mutations (for which one would want to deactivate or destroy the toxic protein).

PARMESAN can also guide genetic modifier screens, as modifier predictions with scores above 10 are 12 times more likely to be proven right than wrong. Modifier screens serve to improve our understanding of gene-gene relationships, which can further guide therapeutic development. For both modifier and drug screens, PARMESAN has the potential to save a tremendous amount of time and resources by identifying the most promising experiments to conduct.

There are three key limitations of PARMESAN that future work will address. 1. The dependence on the availability of literature leads to fewer hypotheses for newly discovered disease genes than for well-known ones. This will be partially addressed by continuing to update PARMESAN’s knowledgebase with new findings, but it may also help to combine the knowledgebase with other publicly available databases such as DrugBank and KEGG. 2. PARMESAN uses abstracts alone, which are far more accessible and easier to parse than full-text articles. Enabling PARMESAN to accurately search open-access, full-text articles will allow it to produce a larger number of promising modifier hypotheses for a gene of interest. 3. The specificity of PARMESAN’s predictions cannot be assessed, since even if all of the hypotheses that it produced were tested, a negative result does not necessarily mean that there is no relationship. The best estimate will therefore come from testing more of PARMESAN’s hypotheses in practice. This will help set a lower bound for specificity, even if an upper bound cannot be ascertained.

Additional future work will include enabling PARMESAN to make additional types of predictions. We have tested its ability to predict drugs and proteins that increase and decrease a given protein’s activity, but it would also be useful to be able to predict treatments that can work downstream of a disease gene, and facilitate the biological processes that the defective protein can no longer facilitate.

In its current state, though, PARMESAN can greatly expedite the literature-search process for known gene-gene and drug-gene relationships, and prioritize drug screens that lead to the discovery of novel therapeutics. This will hopefully lead to effective treatments being brought from the bench to the bedside sooner, and to healthier, happier lives for those who live with severe genetic disorders.

## Methods

### Building a knowledgebase

As it is presented here, PARMESAN uses all PubMed abstracts accessed via PubMed’s FTP site on September 4, 2022. We will present PARMESAN’s extraction process in the context of extracting the relationship described in Monteiro et al, where disrupting *MID1* activity decreases *Huntingtin* levels^15^. This extraction is also displayed diagrammatically in Figure 1. PARMESAN first splits abstracts into sentences, uses PubTator^19^ to determine which words within the abstract are genes, and identifies pairs of genes mentioned in the same sentence. PARMESAN then identifies a Predicate between them. In our example, PARMESAN found the genes “MID1” and “Huntingtin” in the same sentence. The Predicates accepted are listed in Table 1, and the Predicate chosen is the one closest to the second gene mentioned in the sentence. If there is no Predicate, the sentence is not evaluated. The predicate between “MID1” and “Huntingtin” was “reduces”.

PARMESAN then decides whether the sentence is in passive (“Object is affected by Subject”) or active voice (“Subject affects Object”). PARMESAN assumes active voice unless the Predicate is followed by “by” or “in”, indicating passive voice. The sentence is therefore divided into a Subject, Predicate, and Object. The sentence in our example was active voice.

The Subject contains two key elements: The Modifier (the gene in the Subject), and the Action (what is done to the Modifier—such as knocking down). The Action words are in Table 1. If there are multiple possible Actions in the Subject, the one closest to the Modifier is chosen. If two Action words are equidistant from the Modifier, the one before the Modifier is selected. If there is no Action, the sentence is still evaluated. In Monteiro et al, the Subject is “Pharmacological disruption of the MID1/_α_4 interaction”, and the Action is “disruption”.

The Object contains the Target (the gene that the Modifier affects). The Target cannot have the same Entrez gene ID as the Modifier. In Monteiro et al, the Object is “mutant Huntingtin levels in primary neuronal cultures”.

PARMESAN then constructs a summary sentence containing the Action, Modifier, Predicate, and Target. The summary constructed from Monteiro et al is “disruption MID1 reduces Huntingtin”. Since less MID1 activity means less Huntingtin activity, we can infer that more MID1 activity means more Huntingtin activity, which would make MID1 a positive regulator of HTT.

Each relationship, where Gene B increases (positive regulator) or decreases (negative regulator) the level or activity of Gene A, is scored based on the number of articles supporting the relationship:

Let X = the number of articles in which PARMESAN found a summary stating that Gene B positively regulates Gene A.

Let Y = the number of articles in which PARMESAN found a summary stating that Gene B negatively regulates Gene A.

The score D_BA_ is defined per equation (1) below:

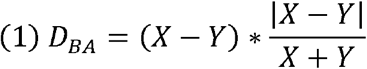

A large positive score indicates that Gene B likely positively regulates Gene A. A large negative score indicates that Gene B likely negatively regulates Gene A. For a large score, there must be proportionally more articles in one direction than in the other.

As an example, in a pre-2010 SNCA modifier search (Supplementary table 11), PARMESAN found 2 articles stating that *GRP78* positively, and 0 stating that it negatively, regulates *SNCA*.

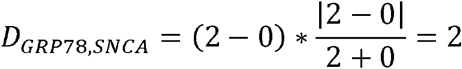

### Accuracy testing

We tested PARMESAN’s ability to construct accurate summaries by manually evaluating 25 random gene-gene and 25 random drug-gene relationships identified before January 1, 2020 (Supplementary tables 1-9). There are three possible outcomes:

1. Correct, where at least one of the supporting abstracts truly confirms the relationship PARMESAN presents, with the same directionality. 2. Misdirected, where at least one of the supporting abstracts truly confirms that the presented modifier affects the presented target, but none of the abstracts agree with the directionality presented by PARMESAN. 3. Incorrect, where none of the supporting abstracts truly indicate that the indicated modifier affects the indicated target.

If the literature indicates that Gene B negatively regulates Gene A, and Gene B has no known effect on Gene C, then a negative-scored Gene B to Gene A relationship would be Correct, a positive-scored Gene B to Gene A relationship would be Misdirected, and a positive- or negative-scored Gene B to Gene C relationship would be Incorrect.

We ran this test four times—once selecting from all of PARMESAN’s relationships, and once selecting from only the relationships whose assigned score (absolute value) was above n, for n = 1, 2, and 3. Because larger values of n limit to relationships with more supporting articles indicating the same directionality, we expected see a higher number of “Correct” findings when n = 3 than when there are no restrictions.

In total, we evaluated 200 relationships: 25 random gene-gene and 25 random drug-gene relationships from PARMESAN whose |D_BA_| > n, for n = 0, 1, 2, and 3.

### Predicting indirect gene-gene and drug-gene relationships

PARMESAN uses known connections to hypothesize on unknown relationships. If Gene C positively regulates Gene B, and Gene B negatively regulates Gene A, then Gene C may negatively regulate Gene A as well.

We score the suspected relationship between C and A based on the quantity and strength of the connections from C to B to A.

Let Q = the set of genes B where C has reported effects on B, and B has reported effects on A.

We define the indirect prediction score I_CA_ per equation (2) below:

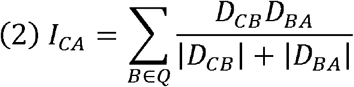

This scoring system favors balanced connections. If one article shows that B positively regulates A, and 200 show that C positively regulates B, that connection increases I_CA_ by 0.995. If 6 articles show that B positively regulates A, and 5 show that C positively regulates B, that connection increases I_CA_ by 2.727. An indirect connection is invalid if either intermediate connection is invalid, and this scoring algorithm downweighs the former case, which is more likely to be invalid than the latter.

### Modeling validation of pre-2010 predictions over time

To model the validation of PARMESAN’s predictions over time, we ran PARMESAN’s gene-gene relationship predictor using only articles published before January 1, 2010. We compared these predictions to PARMESAN’s knowledgebase at timepoints at midnight on January 1 of 2010 and 2020, where PARMESAN’s pre-2010 predictions were compared to the extracted relationships found by that timepoint with an extraction score above 1. The outcome measured was the fraction of pre-2010 predictions validated by each timepoint.

Supplementary tables 10-12 provide an example of this split, showing the full list of predictions of *SNCA* modifiers that PARMESAN made before 2010 (Supplementary table 10), and the full set of extracted *SNCA* modifiers from articles published before (Supplementary table 11) and after (Supplementary table 12) 2010.

### Comparison of predictions with existing knowledge on modifiers and drugs

When comparing relationship predictions to the ground truth, a correct prediction means that PARMESAN’s predicted directionality matched that of the other dataset (both claim that A negatively regulates B). An incorrect prediction means that PARMESAN’s predicted directionality did not match the other dataset (PARMESAN predicts that A positively regulates B, and the dataset states that A negatively regulates B). For every n from 0 to 10, we determine the number of correct and incorrect predictions where the absolute value of the prediction score (I_CA_) is greater than n. Statistical significance between the numbers of correct and incorrect predictions is calculated by one-phase decay least-squares fit with an extra sum-of-squares F test, comparing the decay constant K.

We compared PARMESAN’s drug-gene relationship predictions to the drug-gene relationships presented in DrugBank. To ensure that PARMESAN’s predictions are independent of the information provided by DrugBank, we removed from PARMESAN’s knowledgebase any drug-gene relationships that were taken from an article cited by DrugBank, prior to running the predictive algorithm.

Similarly, we compared PARMESAN’s gene-gene relationship predictions to gene-gene relationships presented by KEGG.

Lastly, we compare PARMESAN gene-gene relationship predictions to screens for modifiers of *ATXN1* and *TAU*. The *ATXN1* screen includes 93 genes whose downregulation leads to reduced *ATXN1* levels (positive regulators identified in Lee et al, which had hits in non-*Drosophila* models)^18^. The TAU screen includes 97 genes whose knockdown decreased (92, positive regulators) or increased (5, negative regulators) endogenous and over-expressed *TAU* protein in a human cell line.

The output of an earlier version of PARMESAN had been compared to the *ATXN1* and *TAU* screens. No modifications were made to any of these screens, with the exception of translating the modifier gene symbols into Entrez Gene IDs. Modifier data were maintained by the Huda Zoghbi lab.

### Statistics

P values comparing accuracies (Supplementary table 9) and the predictions validated over time (Supplementary table 13) are calculated using the binomial distribution Microsoft Excel function, “=1-BINOM.DIST(s,t,p,TRUE))”. For accuracy evaluation, s is the number of accurate relationship extractions with a minimum score of 3, t is the number of relationships evaluated (25), and p is the percent of accurate relationship extractions at a minimum score of 0. For predictions validated over time at a given timepoint, s is the number of validated relationships, t is the number of predicted relationships, and p is the (number of validated / number of predicted) relationships at a minimum score to which we are comparing the validation rate. Each experiment was performed once.

Statistically significant differences between the numbers of correct and incorrect predictions at different minimum scores was calculated by one-phase decay least-squares fit with an extra sum-of-squares F test, comparing the decay constant K. These calculations were done in GraphPad Prism.

## Data availability

The in vitro modifier screens are intellectual property of the Huda Zoghbi lab. Information on the genes in these screens is available upon reasonable request. Modifier extractions from an earlier version of PARMESAN for neurodegenerative disease genes are available at the Neurodegeneration Hub (https://nddb.nrihub.org/). Data tables on drug-gene and gene-gene relationships derived from the DrugBank and KEGG websites are available upon reasonable request that is compliant with the terms of use of the respective organization.

## Code availability

Upon publication, the code used to run PARMESAN and generate a modifier database and set of modifier predictions will be available on a public GitHub repository, at https://github.com/coledeisseroth/PARMESAN.

## Acknowledgements

We thank Roopashri Holehonnur, Nigel Lee, Dongxue Mao, Sasidhar Pasupuleti, Ying-Wooi Wan, Shinya Yamamoto, Megan Mair, Tarik Onur, Juan Botas, and the members of the labs of Huda Zoghbi and Zhandong Liu for their feedback and advice on the improvement and analysis of this tool.

## Author contributions

**CD**: Conceptualization, Methodology, Software, Validation, Formal analysis, Investigation, Data Curation, Writing – Original Draft, Writing – Review & Editing, Visualization. **WSL**: Data Curation, Writing – Review & Editing. **JYK**: Data Curation, Writing – Review & Editing. **HHJ**: Investigation, Data Curation, Writing – Review & Editing. **JW**: Conceptualization, Methodology, Writing – Review & Editing. **HYZ**: Conceptualization, Methodology, Resources, Writing – Review & Editing, Supervision. **ZL**: Conceptualization, Methodology, Resources, Writing – Review & Editing, Supervision, Project administration, Funding acquisition.

## Materials & correspondence

Requests for data from the modifier screens should be addressed to Huda Zoghbi. Other correspondence should be addressed to Zhandong Liu.

