## Supplementary table for "Literature-based predictions of treatments for genetic disease pathology"

**Supplementary materials**

**Supplementary table 1: Evaluation of random gene-gene relationship extractions.** We manually evaluated 25 random gene-gene relationships extracted for accuracy. An extraction is correct if at least one of the source articles confirms the relationship and directionality, and incorrect otherwise. Each relationship description extracted from each article occupies one row. We provide, in order, the PMID of the source article, the modifier, predicate, target, and action extracted from the source article, the Entrez ID of the modifier gene, the Entrez ID of the target gene, the score that PARMESAN gave the relationship, whether the article truly describes the modifier having an effect on the target gene, whether the directionality was correctly extracted, and if the finding was innacurate, a subjective explanation as to why. A mark of “n/a” means that this article was not evaluated, because a different article showed that the relationship was correct. The “Reason” column is a subjective, manually assigned explanation for the cause of an incorrect finding. “Cell type” means the identified gene was used to describe a cell line, not the activity of the gene itself; “Directionality not specified” means that the source article was ambiguous in its description of a possible modifier relationship, and an assumption could not be made; “Incomplete modifier/target” means that a substring of the modifier or target was extracted and interpreted as a different entity; “Negation” means that the relationship was being negated; “Negative action not applicable” means that the negative action extracted should not have been used to reverse the directionality of the extracted relationship; “Negative action [on target]” means that a word that should have been used to reverse the directionality of the extracted relationship was not detected; “No modifier relationship described” means that there was no relationship that should have been extractable; “Not a gene” means that one of the entities extracted that should have been a gene was not; “Overridden” means that this particular extraction disagrees with PARMESAN’s overall assessment of the directionality of the modifier relationship—we mark this as correct if the source abstract agrees with PARMESAN’s overall modifier score, and misdirected otherwise; “Reversed” means that PARMESAN extracted a relationship from A to B, when the true relationship from the abstract was from B to A; “Wrong modifier/predicate/target” means that the specified element should have been taken from a different word/phrase in the abstract for the extraction to be accurate.

**Supplementary table 2: Evaluation of random gene-gene relationship extractions, minimum score of 1.** We manually evaluated 25 random gene-gene relationships (among those to which PARMESAN assigned a score of at least 1) extracted for accuracy. This table has the same format as Supplementary Table 1.

**Supplementary table 3: Evaluation of random gene-gene relationship extractions, minimum score of 2.** We manually evaluated 25 random gene-gene relationships (among those to which PARMESAN assigned a score of at least 2) extracted for accuracy. This table has the same format as Supplementary Table 1.

**Supplementary table 4: Evaluation of random gene-gene relationship extractions, minimum score of 3.** We manually evaluated 25 random gene-gene relationships (among those to which PARMESAN assigned a score of at least 3) extracted for accuracy. This table has the same format as Supplementary Table 1.

**Supplementary table 5: Evaluation of random drug-gene relationship extractions.** We manually evaluated 25 random drug-gene relationships extracted for accuracy. This table has the same format as Supplementary Table 1.

**Supplementary table 6: Evaluation of random drug-gene relationship extractions, minimum score of 1.** We manually evaluated 25 random drug-gene relationships (among those to which PARMESAN assigned a score of at least 1) extracted for accuracy. This table has the same format as Supplementary Table 1.

**Supplementary table 7: Evaluation of random drug-gene relationship extractions, minimum score of 2.** We manually evaluated 25 random drug-gene relationships (among those to which PARMESAN assigned a score of at least 2) extracted for accuracy. This table has the same format as Supplementary Table 1.

**Supplementary table 8: Evaluation of random drug-gene relationship extractions, minimum score of 3.** We manually evaluated 25 random drug-gene relationships (among those to which PARMESAN assigned a score of at least 3) extracted for accuracy. This table has the same format as Supplementary Table 1.

**Supplementary table 9: Accuracy test results.** Here we provide the numbers of correct, misdirected, and incorrect relationship extractions from Supplementary Tables 1-8, and the calculated binomial distribution p values comparing the accuracy at a minimum score of 3 to that at a minimum score of 0.

**Supplementary table 10: Pre-2010 *SNCA* predictions**. We present the genetic modifier predictions made by PARMESAN for *SNCA*, using only articles published before January 1, 2010. Each entry provides a two-step hypothesis, where Gene B modifies Gene A (Step 1), and Gene A modifies *SNCA* (Step 2). The first column, “Step 1 PMID”, is the PubMed ID of the article showing that Gene B modifies Gene A. The second column, “Step 2 PMID”, is the PubMed ID of the article showing that Gene A modifies *SNCA*. The third column, “Predicted modifier (Entrez ID)” is the Entrez ID of Gene B. The fourth column, “Intermediate modifier (Entrez ID)”, is the Entrez ID of Gene A. The fifth column, “Step 1 score” is the score that PARMESAN gave the relationship between Gene B and Gene A (D_BA_). The fifth column, Step 2 score, is the score that PARMESAN gave the relationship between Gene A and *SNCA* (D_A,_*_SNCA_*). The sixth column, “Modifier score”, is the score that PARMESAN gave the predicted relationship between Gene B and *SNCA* (I_B,_*_SNCA_*). The seventh column, “Hypothesis”, is an automatically generated sentence describing the hypothesis formed by the two given articles. This is only one example for one gene, and the code needed to generate hypotheses for other genes will be made available upon publication.

**Supplementary table 11: Pre-2010 *SNCA* modifiers.** We present the genetic modifiers identified by PARMESAN for *SNCA*, using only articles published before January 1, 2010. The first column is the PubMed ID of the article describing the relationship. The second column is the Entrez ID of the modifier. The third column is the score that PARMESAN gave the relationship between the modifier and *SNCA* (D_A,_*_SNCA_*). The fourth column is an automatically generated sentence describing the relationship extracted from the article.

**Supplementary table 12: Pre-2020 *SNCA* modifiers.** We present the genetic modifiers identified by PARMESAN for *SNCA*, using only articles published before January 1, 2020. The first column is the PubMed ID of the article describing the relationship. The second column is the Entrez ID of the modifier. The third column is the score that PARMESAN gave the relationship between the modifier and *SNCA* (D_A,_*_SNCA_*). The fourth column is an automatically generated sentence describing the relationship extracted from the article.

**Supplementary table 13: Predictions validated over time.** We limited PARMESAN’s dataset to articles from before 2010, and observed the percentage of its gene-gene and drug-gene relationship predictions (with different minimum prediction confidence scores) for that were validated before 2010 and before 2020. “Validated” means consistent with the PARMESAN’s knowledgebase of direct relationships (with an extraction confidence score of at least 1) at the given timepoint. We display the minimum prediction confidence score, the number of predictions made, the number and fraction of pre-2010 predictions that were consistent with PARMESAN’s knowledgebase before 2010, the number and fraction of pre-2010 predictions that were consistent with PARMESAN’s knowledgebase before 2020, and the increase in the fraction of predictions validated from 2010 to 2020.

**Supplementary table 14: DrugBank comparison.** We compared PARMESAN’s drug-gene relationship predictions (excluding the articles cited by DrugBank) to the drug-gene relationships presented by DrugBank. “Correct” means that PARMESAN’s prediction matched the directionality of the relationship displayed in DrugBank (whether Drug A increases or decreases the activity of Gene B), and “Incorrect” means that PARMESAN’s prediction had the opposite directionality.

**Supplementary table 15: KEGG comparison.** We compared PARMESAN’s gene-gene relationship predictions to the gene-gene relationships presented by KEGG. “Correct” means that PARMESAN’s prediction matched the directionality of the relationship displayed in KEGG (whether Gene A increases or decreases the activity of Gene B), and “Incorrect” means that PARMESAN’s prediction had the opposite directionality.

**Supplementary table 16: Modifier screen comparison.** We compared PARMESAN’s gene-gene relationship predictions to *in vitro* screens for modifiers of *ATXN1* and *TAU*. “Correct” means that PARMESAN’s prediction matched the directionality of the relationship identified in the screen, and “Incorrect” means that PARMESAN’s prediction had the opposite directionality.
